# Nucleosome-binding by TP53, TP63, and TP73 is determined by the composition, accessibility, and helical orientation of their binding sites

**DOI:** 10.1101/2024.05.03.592419

**Authors:** Patrick D. Wilson, Xinyang Yu, Michael J. Buck

## Abstract

The p53 family of transcription factors plays key roles in driving development and combating cancer by regulating gene expression. TP53, TP63, and TP73—the three members of the p53 family—regulate gene expression by binding to their DNA binding sites, many of which are situated within nucleosomes. To thoroughly examine the nucleosome-binding abilities of the p53 family, we used Pioneer-seq, a technique that assesses a transcription factor’s binding affinity to its DNA binding sites at all possible positions within the nucleosome core particle. Using Pioneer-seq, we analyzed the binding affinity of TP53, TP63, and TP73 to 10 p53-family binding sites across the nucleosome core particle. We found that the affinity of TP53, TP63, and TP73 for nucleosomes was largely determined by the positioning of p53-family binding sites within nucleosomes; p53-family members bind strongly to the more accessible edges of nucleosomes but weakly to the less accessible centers of nucleosomes. We also found that the DNA-helical orientation of p53-family binding sites within nucleosomal DNA impacted the nucleosome-binding affinity of p53-family members. The composition of their binding sites also impacted each p53-family member’s nucleosome-binding affinities only when the binding site was located in an accessible location. Taken together, our results show that the accessibility, composition, and helical orientation of p53-family binding sites collectively determine the nucleosome-binding affinities of TP53, TP63, and TP73. These findings help explain the rules underlying p53-family-nucleosome binding and thus provide requisite insight into how we may better control gene-expression changes involved in development and tumor suppression.

## INTRODUCTION

The p53 family of transcription factors induces the expression of genes that determine the survival, proliferation, and differentiation of cells. In this family are three members: TP53, TP63, and TP73. TP53 has a well-known role in tumor suppression and is mutated in more than 50% of human cancers (Joerger and Fersht 2007; Baugh et al. 2018; Marei et al. 2021). Once activated in response to cell-stress signals, TP53 induces the expression of genes involved in cell-cycle arrest and apoptosis (Nguyen et al. 2018). TP63 and TP73 also induce the expression of some of these TP53-regulated genes (Moll and Slade 2004), though both transcription factors play more of a prominent role in the induction of developmental genes. TP63 induces the expression of genes involved in the proliferation, differentiation, and adhesion of epithelial cells (Kouwenhoven et al. 2015; Sethi et al. 2015; Boughner et al. 2018), while TP73 induces the expression of genes involved in neuronal development, multiciliogenesis, germ-cell maturation, and angiogenesis (Nemajerova and Moll 2019; Logotheti et al. 2021). The gene-regulatory complexity of the p53-family is further increased by the existence of multiple isoforms of each p53-family member that have overlapping and sometimes opposing functions (Wei et al. 2012).

Acting in their capacity as transcription factors, p53-family members induce gene expression through their binding to specific DNA sequences, also known as ‘binding sites.’ TP53 specifically binds to a 20-base-pair binding site comprising two repeats of RRRCWWGYYY (where R = A or G; W = A or T; Y = C or T) separated by 0-13 base pairs (el-Deiry et al. 1992; Funk et al. 1992)— though TP53 also binds to a large number of sequences that deviate from this binding-site pattern (Tomso et al. 2005; Veprintsev and Fersht 2008). TP63 and TP73 can bind to the same binding sites as TP53 (Osada et al. 2005; Lokshin et al. 2007; Schavolt and Pietenpol 2007), owing to the high percentage of sequence identity between their DNA-binding domains (Levrero et al. 2000). Each p53-family member binds to these 20-base-pair binding sites (hereafter termed ‘p53-family binding sites’) as a tetramer, specifically a dimer of dimers that are each bound to an RRRCWWGYYY repeat (Brandt et al. 2009).

Eukaryotic DNA is organized into chromatin, which is made up of nucleosomes (Cooper 2000). A nucleosome comprises an octamer of histone proteins (2 copies each of H2A, H2B, H3, and H4) around which ~146 base pairs of DNA are coiled (Luger et al. 1997). Although the DNA coiled within nucleosomes is typically inaccessible to transcription-factor binding (Zhu et al. 2018), TP53 is able to bind to instances of its binding sites within nucleosomes (Espinosa and Emerson 2001; Lidor Nili et al. 2010; Sammons et al. 2015), specifically when positioned near the nucleosome edges (Laptenko et al. 2011; Yu and Buck 2019; Nishimura et al. 2020). TP63 is also known to bind to instances of its binding sites within nucleosomes (Sethi et al. 2014; Yu et al. 2021), though TP73 has not yet been tested in any nucleosome-binding studies. Despite the remarkable progress in defining the capabilities of the TP53 family to bind to nucleosomes, little is known of the underlying intricacies of p53-family-nucleosome binding.

In this study, we measured the relative binding affinity of each p53-family member to 10 different p53-family binding sites when positioned at every base pair of the Widom-601 nucleosome. This comprehensive dataset allowed us to reveal the interplay between nucleosomal context and p53-family binding. We found that p53-family-nucleosome binding is collectively shaped by the DNA-sequence composition, nucleosomal accessibility, and DNA-helical orientation of p53-family binding sites within the nucleosome. Our findings shed light on the underlying principles of p53-family-nucleosome binding, offering insight into how we can effectively modulate p53-family-mediated gene-expression changes in tumor suppression and development.

## RESULTS

### Pioneer-seq reveals the nucleosome-binding preferences of TP53, TP63, and TP73

To comprehensively examine the nucleosome-binding abilities of the p53 family, we developed Pioneer-seq, which is an extension of low-throughput nucleosome-binding assays (Yu and Buck 2019). Pioneer-seq is a high-throughput competitive nucleosome-binding assay for gauging the binding affinities of transcription factors to their transcription-factor binding sites at every base-pair position of the nucleosome core particle and at several positions of the linker DNA. Pioneer-seq is similar to SeEN-seq, which has been successfully used to identify TF-bound nucleosomes for cryo-EM (Michael et al. 2020; Michael et al. 2023). SeEN-seq has been limited to a small nucleosome library, less than 200 sequences with a single TFBS variant. Outlined in Fig. 1a, the Pioneer-seq protocol starts with the in-silico-design of a library of Widom-601 nucleosome-positioning sequences that each contain an individual transcription-factor binding site at a unique position of the nucleosome core particle or linker DNA. This DNA library is then reconstituted into a nucleosome library and incubated with the transcription factor of interest. Stable transcription-factor-nucleosome complexes are then isolated via gel-shift assay (Supplemental Fig. 1) and DNA-sequenced. The resulting sequencing data is then analyzed to gauge the binding affinity of the transcription factor of interest to its binding site at each nucleosomal position (**Equation 1**). For this study, we performed Pioneer-seq experiments with TP53, TP63, and TP73 using a nucleosome library that contained 10 p53-family binding sites (**Supplementary Table 1**) individually positioned at every base pair of the Widom-601 nucleosome core particle and at several positions of the linker DNA.

**Figure 1.**
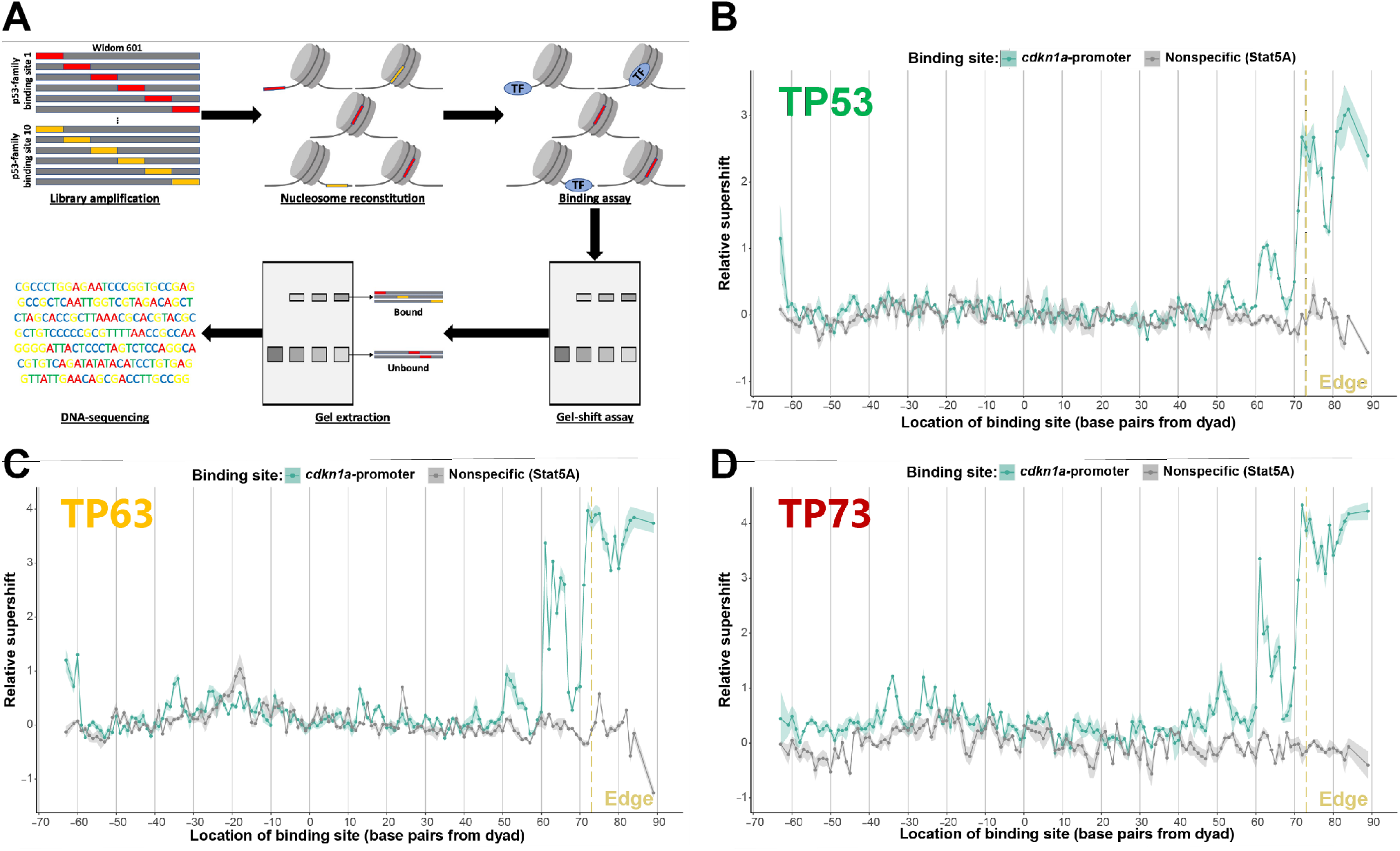
Pioneer-seq maps TP53-family-nucleosome binding. (A) Outline of Pioneer-seq. A library of 217 base-pair nucleosome-positioning sequences, which are based on the Widom-601 nucleosome-positioning sequence, and which have a TP53-family binding site in one of the 146 base-pair positions of the nucleosome core particle, is reconstituted into nucleosomes and then incubated with TP53, TP63, or TP73. Then TP53-family-bound library nucleosomes are separated from unbound library nucleosomes via gel-shift assay, and the gel bands corresponding to the TP53-family-bound and - unbound library nucleosomes are then extracted, purified, and sequenced. (B-D) TP53, TP63, or TP73 binding to a cdkn1a-promoter TP53-family binding site at all 146 base-pair positions of the nucleosome core particle and at several positions of the linker DNA. TP53-family binding to the cdkn1a-promoter binding site is measured relative to TP53-family binding to nonspecific (non-TP53-family) binding sites and represented by relative-supershift values (Equation 1). ‘Edge’ = the right end of the 146 base-pair nucleosome core particle. Shading = SEM. n = 3.

We first examined how strongly each p53-family member binds to a p53-family binding site from the *cdkn1a* promoter at different positions of the Widom-601 nucleosome. The predominant pattern of p53-family-nucleosome binding was that the affinity each p53-family member has for their *cdkn1a*-promoter binding site is dependent on how far it is positioned from the dyad of the nucleosome core particle. Notably, though, there were exceptions to this pattern—e.g.: TP53, TP63, and TP73 bind more strongly to their *cdk1na*-promoter binding site when it is 65 base pairs away from the dyad than when it is 68 base pairs away (**Fig. 1b-c**). Our findings reveal the significant extent to which p53-family binding is influenced by the precise position of p53-family binding sites within nucleosomes.

### Sequence composition of binding sites modulates TP53-, TP63-, and TP73-nucleosome binding affinity

Having established where p53-family members could bind to their *cdkn1a*-promoter binding site within the Widom-601 nucleosome, we then sought to identify the specific factors that underlie where p53-family members can bind nucleosomes. It seemed likely that one such factor was the sequence-composition of p53-family binding sites, given the significant impact the sequence-composition of TP53 binding sites has on TP53-naked-DNA binding affinity. This prompted us to investigate how mutations in p53-family binding sites impact the ability of p53-family members to bind to nucleosomes. We measured how well TP53, TP63, and TP73 bind to unmutated and mutated versions of a high-affinity p53-family binding site (**Fig. 2a**) across the Widom-601 nucleosome core particle and linker DNA. At nearly all of the Widom-601 nucleosome core particle and linker positions where p53-family members could bind to their high-affinity binding site, p53-family members bind with a lower affinity to the once-mutated version of this binding site, and bind with an even lower affinity to the twice-mutated version of this binding site (**Fig. 2b, d, f**). To validate these Pioneer-seq results, we conducted traditional nucleosome-binding assays using two selected nucleosomes from the Pioneer-seq library: one with the high-affinity binding site 70 base pairs from the dyad and the other with the once-mutated binding site at the same distance (**Fig. 2b**). These traditional nucleosome-binding assays confirmed that altering a single base pair of a p53-family binding site significantly decreased the affinity of TP53, TP63, and TP73 for nucleosomes (**Fig. 2c, e, g**).

**Figure 2.**
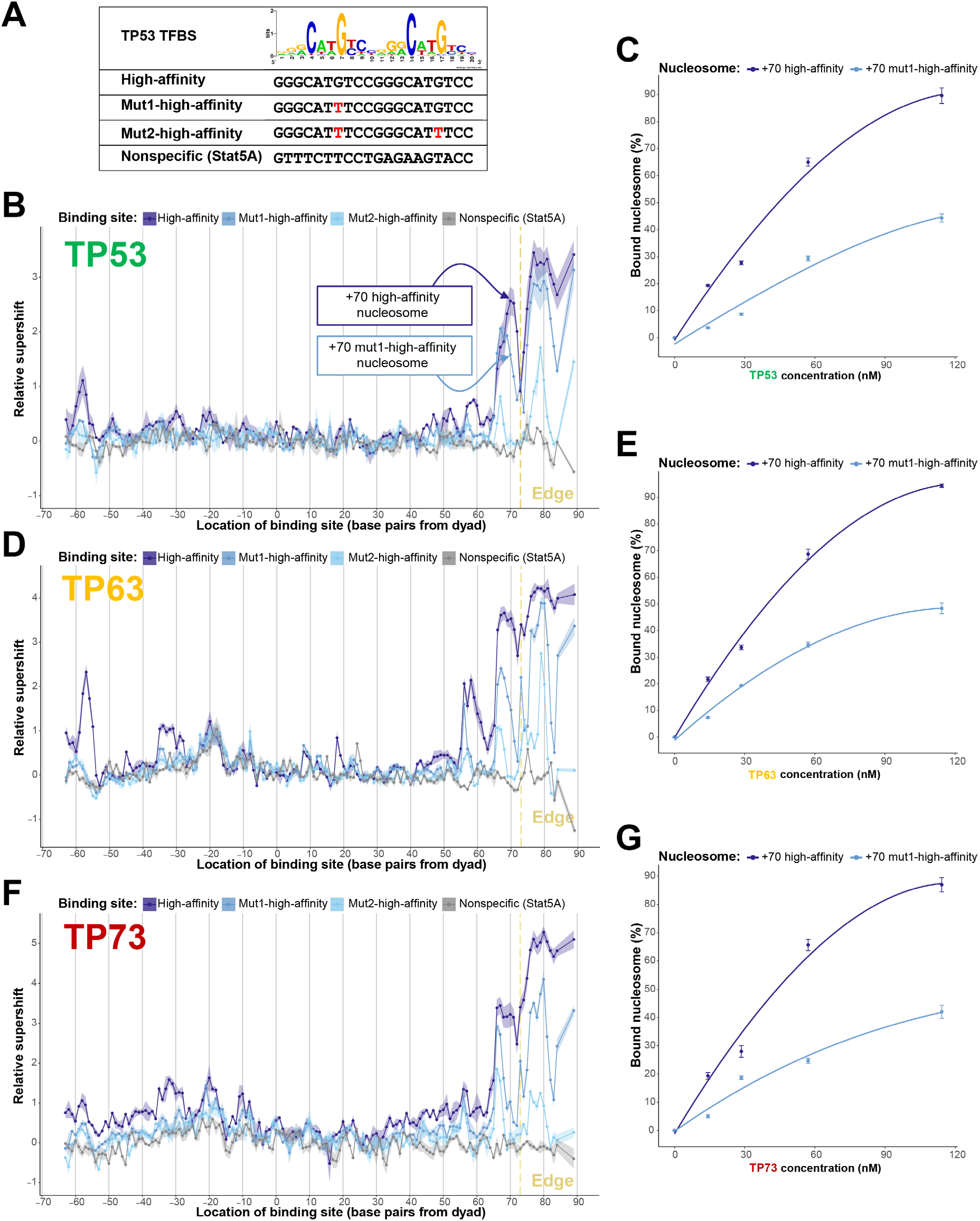
Binding-site composition modulates TP53-family-nucleosome binding affinity. (A) A TP53 sequence logo displayed above the following binding sites included in the nucleosome library: High-affinity binding site, based on the TP53 sequence logo; Mut1-high-affinity binding site, also based on the TP53 sequence logo but with one of the most highly conserved bases mutated; Mut2-high-affinity binding site, also based on the TP53 sequence logo but with two of the most highly conserved bases mutated; a Stat5A binding site nonspecifically targeted by TP53-family members. (B, D, F) TP53, TP63, or TP73 binding to the four binding sites in (A) at all 146 base-pair positions of the nucleosome core particle and at several positions of the linker DNA. TP53-family binding to each binding site is measured relative to TP53-family binding to nonspecific (non-TP53-family) binding sites and represented by relative-supershift values (Equation 1). ‘Edge’ = the right end of the 146-base-pair nucleosome core particle. Shading = SEM. n = 3. (C, E, F) Assessing the binding affinity between TP753, TP63, or TP73 and the +70 high-affinity nucleosome or the +70 mut1-high-affinity nucleosome—both of which nucleosomes are indicated in (B). Bound nucleosome (%) was calculated via gel-shift assays featuring Cy5-labelled +70 high-affinity or +70 mut1-high-affinity nucleosomes.

### Binding-site accessibility is a key determinant of TP53-family-nucleosome binding

From our Pioneer-seq data, it appeared that the proximity of p53-family binding sites to the dyad of the nucleosome core particle was inversely correlated with the affinity of p53-family members for nucleosomes (**Fig. 1b-d**; **Fig. 2b, d, f**). We questioned whether this was because there was a relationship between p53-family-nucleosome binding affinity and binding-site accessibility. To address this question, we digested the nucleosomes in the Pioneer-seq library using micrococcal nuclease (MNase), which is a nonspecific endo- and exonuclease sensitive to the accessibility of nucleosomal DNA (Tsompana and Buck 2014). The susceptibility of library nucleosomes to MNase digestion mirrored the structural organization of the nucleosome core particle, with the tightly wrapped dyad being most protected from MNase digestion and the more loosely wrapped edges being least protected from digestion (**Fig. 3a**). Then we analyzed the MNase-digested Pioneer-seq library to measure how accessible the *cdkn1a*-promoter and high-affinity p53-family binding sites are at each base-pair position of the Widom-601 nucleosome. Our analysis led us to find a strong correlation between *cdkn1a*-promoter- and high-affinity-binding-site accessibility and p53-family-nucleosome binding affinity. Further analysis of the MNase-digested Pioneer-seq library revealed that the accessibility of other p53-family binding sites—i.e., binding sites from the promoters of the *puma, ppn1*, and *chm4pc* genes—also correlated with p53-family-nucleosome binding affinity (**Fig. 3b-d**). This correlation between binding-site accessibility and p53-family-nucleosome binding appeared dependent on the affinity of the p53-family for the respective binding site, since this correlation was weaker for the once-mutated and twice-mutated binding sites than for the high-affinity binding site (**Supplementary Fig. 2**).

**Figure 3.**
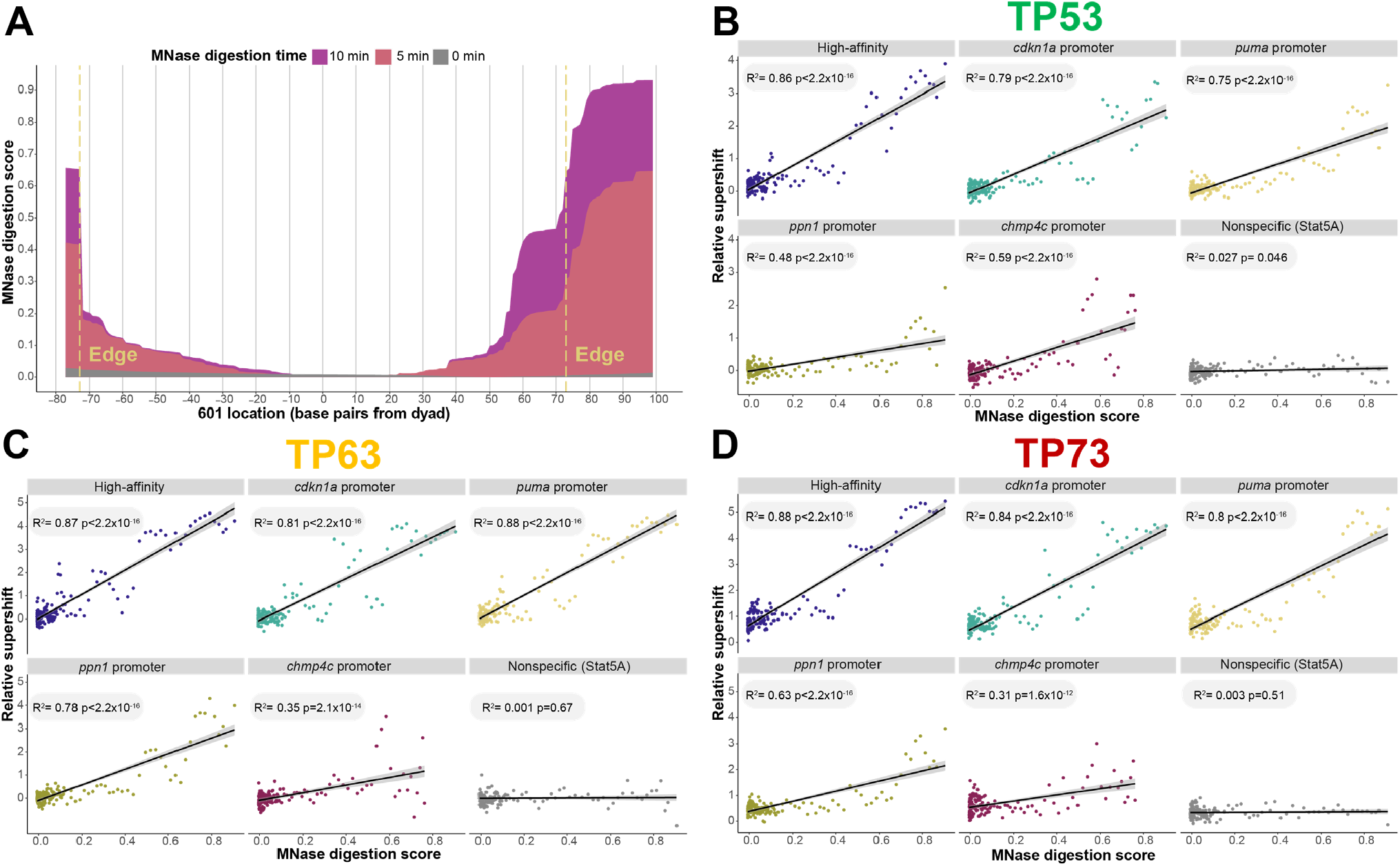
Binding-site accessibility is a key determinant of TP53-family-nucleosome binding. (A) Measuring library-nucleosome accessibility via MNase digestion. MNase was incubated with the nucleosome library for 0, 5, or 10 minutes, followed by DNA-sequencing to measure MNase digestion at base-pair resolution, which is represented by MNase-digestion scores. For each base pair in the library nucleosomes, MNase-digestion scores were computed as the ratio of the number of reads covering that base pair to the total number of reads mapped to the respective library nucleosome. ‘Edge’ = the left or right end of the canonical 146-base-pair nucleosome core particle. (B-D) Correlating the binding strength of TP53, TP63, or TP73 to five TP53-family binding sites and one nonspecific (Stat5A) binding site with the nucleosomal accessibility of these binding sites. TP53-family binding to each binding site is measured relative to TP53-family binding to nonspecific (non-TP53-family) binding sites and represented by relative-supershift values (Equation 1). Nucleosomal accessibility of binding sites is measured by averaging the MNase-digestion scores of each base pair in the binding site. Shading around regression lines represents 95% confidence interval.

### Binding-site helical orientation in DNA impacts p53-family-nucleosome binding

Although the composition and accessibility of binding sites are together strong determinants of p53-family-nucleosome binding, they still do not fully explain the patterns of p53-family-nucleosome binding we observed. See for instance how TP53 binds with a greater affinity to its high-affinity binding site when positioned 70 base pairs away from the dyad of the Widom-601 nucleosome core particle than when positioned 73 base pairs away (**Fig. 4a-b**), even though this latter position is more accessible (**Fig. 3a**). This discrepancy suggested to us the involvement of an unrecognized factor in regulating p53-family-nucleosome binding. To potentially identify this unrecognized factor, we structurally modelled a Widom-601 nucleosome with the high-affinity p53-family binding site 70 base pairs away from the dyad of the nucleosome core particle (the ‘+70 high-affinity nucleosome’) and a Widom-601 nucleosome with this high-affinity binding site 73 base pairs away (the ‘+73 high-affinity nucleosome’) (**Fig. 4a**). In the strongly TP53-bound +70 high-affinity nucleosome, the most highly conserved bases in the high-affinity p53-family binding site—the CATGs (**Fig. 2a**)—are positioned in the major grooves of the nucleosomal DNA; whereas in the weakly TP53-bound +73 high-affinity nucleosome, these CATGs are positioned in the minor grooves (**Fig. 4c**). It appeared from this observation that the helical orientation of p53-family binding sites impacts p53-family nucleosome binding.

**Figure 4.**
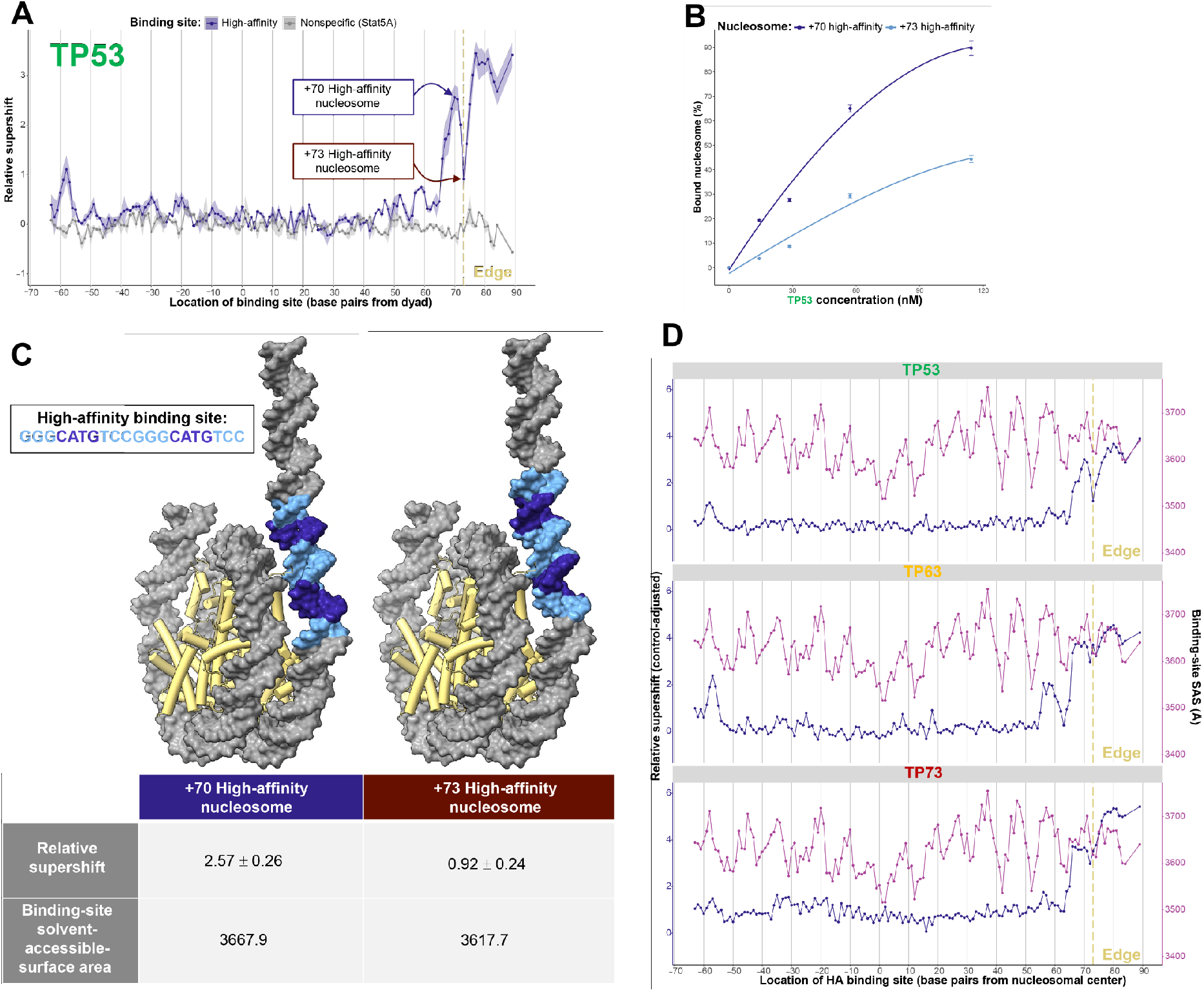
Binding-site helical orientation in nucleosomal DNA impacts TP53-family-nucleosome binding. (A) TP53 binding to a high-affinity (HA) TP53-family binding site at all 146 base-pair positions of the nucleosome core particle and at several positions of the linker DNA. TP53-family binding to the HA binding site is measured relative to TP53-family binding to nonspecific (non-TP53-family) binding sites and represented by relative-supershift values (Equation 1). ‘Edge’ = the right end of the 146-base-pair nucleosome core particle. Shading = SEM. n = 3. (B) Comparing the binding affinity between TP53 and the +70 high-affinity nucleosome or the +73 high-affinity nucleosome—both of which nucleosomes are indicated in (A). Bound nucleosome (%) was calculated via gel-shift assays featuring Cy5-labelled +70 high-affinity or +73 high-affinity nucleosomes. (C) Illustrative models of the +70 and +73 high-affinity nucleosomes indicated in (A). The high-affinity binding site in each nucleosome is highlighted in blue, with the most conserved bases (see Fig. 2a) highlighted in dark blue. The TP53-relative-supershift value corresponding to each nucleosome is shown. Also shown is the solvent-accessible-surface area of the conserved CATG bases of the high-affinity binding site in each nucleosome. (D) TP53, TP63, or TP73 binding to the HA binding site at all 146 base-pair positions of the nucleosome core particle and at several positions of the linker DNA is plotted in blue as ‘control-adjusted’ relative-supershift values (i.e., relative-supershift values of nonspecific-binding-site-containing nucleosomes subtracted from relative-supershift values of HA-binding-site-containing nucleosomes). Solvent-accessible-surface area (SAS) of the conserved CATG bases of the high-affinity binding site is plotted in purple. ‘Edge’ = the right end of the 146-base-pair nucleosome core particle.

To further investigate the role of the helical orientation of binding sites in p53-family-nucleosome binding, we used ChimeraX to measure the solvent-accessible surface area (SASA) of the CATGs of the high-affinity p53-family binding site. These measurements provided insights into the relative exposure of the binding-site CATGs to the solvent environment, enabling us to assess their potential accessibility to p53-family binding. We found that the binding-site CATGs in the strongly TP53-bound +70 high-affinity nucleosome were more exposed than the binding-site CATGs in the weakly TP53-bound +73 high-affinity nucleosome (**Fig. 4c**). Then, to comprehensively assess the role of the helical orientation of binding sites in p53-family-nucleosome binding, we extended our SASA measurements to encompass every library nucleosome that has a high-affinity p53-family binding site. At nucleosomal positions with high binding-site accessibility but unexpectedly low p53-family binding, we found that the binding-site CATGs have consistently lower SASA values compared to positions with higher p53-family binding (**Fig. 4d**).

### p53-family-nucleosome binding is a function of the composition, accessibility, and helical orientation of p53-family binding sites

Our analysis revealed three factors underlying p53-family-nucleosome binding: binding-site composition, binding-site accessibility, and binding-site DNA-helical orientation. We wished to see whether these factors have a synergistic effect on p53-family-nucleosome binding, so we performed stepwise multiple-regression analysis. This statistical approach iteratively selected the most relevant variables from a larger pool of candidate variables based on their statistical significance in explaining our observed patterns of p53-family-nucleosome binding. The initial candidate variables comprised not only scores reflecting each of the three factors underlying p53-family-nucleosome binding but also all pairwise interactions between them. From this stepwise multiple-regression analysis, we derived a model for predicting p53-family-nucleosome binding (**Fig. 5a**).

**Figure 5.**
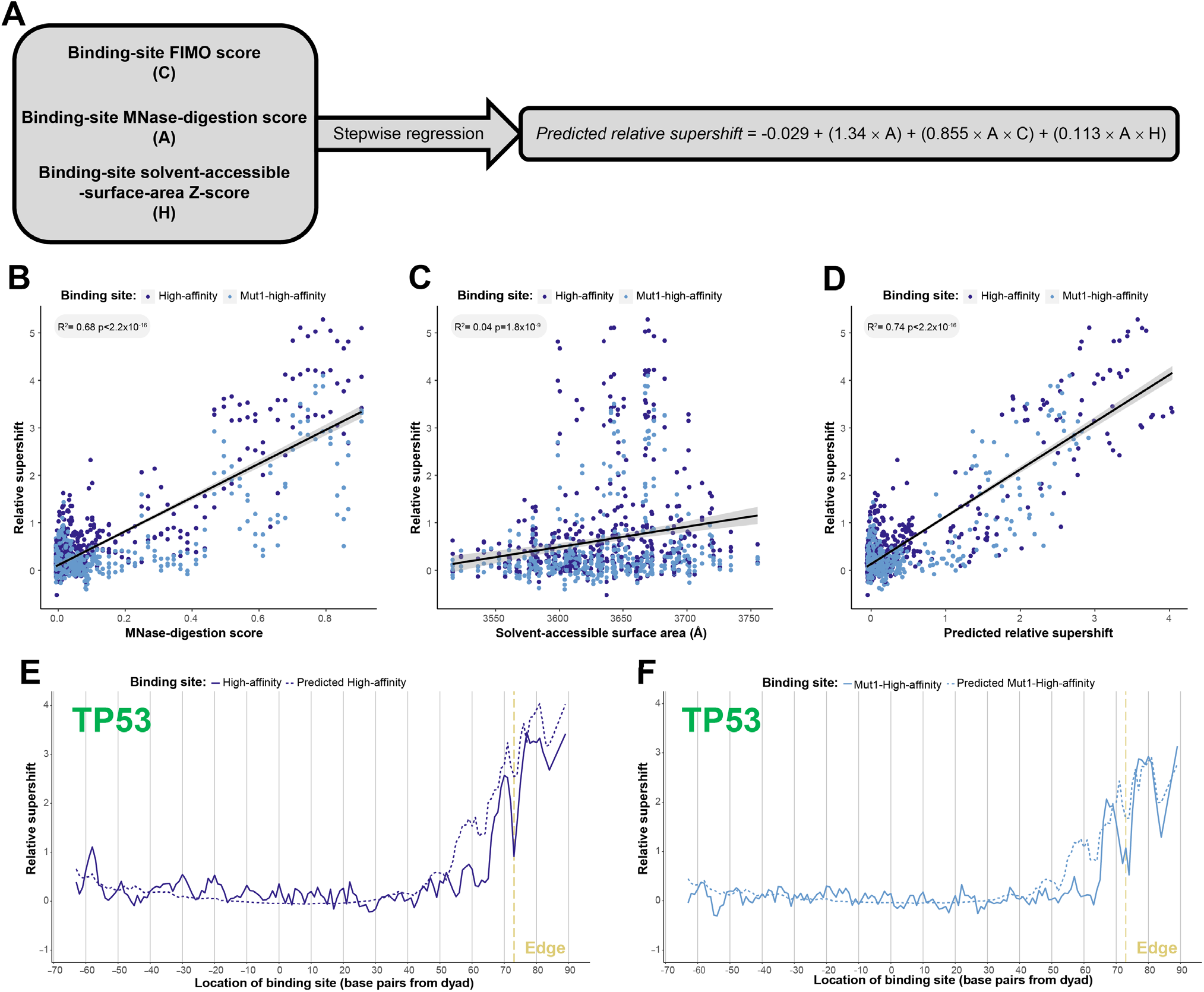
TP53-family-nucleosome binding is collectively determined by the composition, accessibility, and helical orientation of TP53-family binding sites. (A) Outline of the process for generating a multiple regression model predicting TP53-family-nucleosome binding. Three independent variables—binding-site FIMO scores, binding-site MNase-digestion scores, and binding-site solvent-accessible-surface-area z-scores—were inputs of a stepwise regression model. The resulting equation illustrates the predictive relationship between these variables and TP53-family-nucleosome binding. (B-D) Correlating the relative-supershift values (Equation 1) of TP53, TP63, and TP73 to two TP53-family binding sites with either the MNase-digestion score of these binding sites, the solvent-accessible surface area of the CATGs of these binding sites, or the relative-supershift values of TP53, TP63, and TP73 to these binding sites predicted by the multiple regression model outlined in (A). (E, F) Comparing actual relative-supershift values of TP53 to the high-affinity TP53-family binding site or mut1-high-affinity TP53-family binding site with model-predicted relative-supershift values.

To see whether binding-site composition, accessibility, and orientation have a synergistic effect on p53-family-nucleosome binding, we compared the ability of our predictive model to accurately explain p53-family-nucleosome binding to the explanatory capacity of either binding-site accessibility or helical orientation alone. While binding-site accessibility alone explained 68% of the variance observed in p53-family-nucleosome binding (**Fig. 5b**), and binding-site helical orientation alone explained only 4% (**Fig. 5c**), our multivariate model explained a notable 74% of the variance observed in p53-family-nucleosome binding (**Fig. 5d**). This improvement in explanatory power demonstrates that binding-site composition, orientation, and accessibility are not acting independently but are interconnected and work together to the determine the binding affinity of p53-family members for their nucleosomal binding sites.

## DISCUSSION

This study explored the interplay between the p53 family of transcription factors, their binding sites, and the nucleosome, revealing three critical factors influencing p53-family-nucleosome binding: (1) the DNA-sequence composition of p53-family binding sites, (2) the accessibility of these binding sites in the nucleosome, and (3) the helical orientation of these binding sites in the nucleosome. These factors collectively shape the binding landscape of the p53 family to nucleosomes. This suggests that p53-family binding is fine-tuned by the nucleosomal context and DNA-sequence composition of p53-family binding sites—e.g., a less favorable p53-family binding site could be bound strongly if positioned in a more accessible region and oriented in a more exposed orientation in the nucleosome.

Building on previous findings that the DNA-sequence composition of p53-family binding sites significantly impacts the affinity of p53-family members for naked DNA (Veprintsev and Fersht 2008; Brandt et al. 2009), our findings show that the composition of p53-family binding sites significantly impacts the affinity of p53-family members for nucleosomal DNA. Our findings also show that the position of p53-family binding sites within nucleosomes strongly influences p53-family binding. The regions of nucleosomes that are bound most strongly by p53-family members—the edges—are the most accessible regions of nucleosomes, which aligns with the prevailing model of dynamic partial DNA-unwrapping from histones at the edges of nucleosomes (Kitayner et al. 2006; Beno et al. 2011; Jordan et al. 2012). Our findings also show that the helical orientation of p53-family binding sites in nucleosomal DNA has a considerable effect on the p53-family binding. This finding lends further credence to the proposed hypothesis that the helical orientation of nucleosomal TP53 binding sites impacts p53-nucleosome binding affinity, which in turn may impact the differential induction patterns of p53-regulated genes (Freewoman et al. 2021).

From previous studies, it had already emerged that the affinity TP53 and TP63 have for nucleosomes is dependent on the positioning of their binding sites (Yu and Buck 2019; Nishimura et al. 2020; Yu et al. 2021). This led us to expect p53-family-nucleosome binding affinity to simply increase as p53-family binding sites were positioned further from the dyad of the nucleosome. While our initial expectation mostly held true, there were notable exceptions of weak p53-family-nucleosome binding despite far positioning of the p53-family binding site from the dyad. The high-throughput nature of our Pioneer-seq assay enabled us to detect these nucleosome-binding deviations, which were less obvious in previous studies examining p53-family binding at only a few nucleosomal positions. Detecting these deviations in p53-family-nucleosome binding allowed us to identify the helical orientation of p53-family binding sites as an important factor underlying p53-family-nucleosome binding.

While TP53’s affinity for specific DNA sequences (i.e., its binding sites) in nucleosomes is well-established, our previous work showed that TP53 also has a considerable affinity for nonspecific DNA sequences in nucleosomes, mediated by its C-terminal domain (Yu and Buck 2019). These nonspecific TP53-nucleosomal-DNA binding events were illustrated recently in a set of cryo-electron-microscopy experiments, revealing that the C-terminal domain of TP53 binds to the dyad and linker-DNA of nucleosomes (Nishimura et al. 2022). In the present study, however, we were unable to distinguish between binding-site-specific and nonspecific p53-family-nucleosome binding, on account of Pioneer-seq being a competitive nucleosome-binding assay. Pioneer-seq measures the overall affinity of a transcription factor for a nucleosome and thus cannot resolve the distinct contributions of binding-site-specific and -nonspecific binding to the measured affinities of p53-family members for nucleosomes. To address this limitation, it would be beneficial for future studies to investigate the contributions of individual residues within the domains of p53-family members to nucleosome-binding affinity and specificity. This could be achieved through mutagenesis studies that introduce specific amino-acid substitutions in the p53-family proteins and then assess their ability to bind to nucleosomes. By systematically mutating key p53-family residues and evaluating the resulting effects on nucleosome-binding affinity and specificity, a more comprehensive view of the molecular mechanisms underlying p53-family-nucleosome binding could be achieved.

In this study, we explored the intricate interplay between the p53 family of transcription factors, their binding sites, and the nucleosome to show that p53-family-nuclesome binding is collectively shaped by the DNA-sequence composition of p53-family binding sites, the accessibility of these binding sites in the nucleosome, and the helical orientation of these binding sites in the nucleosome. This has significant implications for comprehending how p53-family members regulate gene expression in various cellular processes, including development and tumor suppression. By identifying the factors underlying p53-family-nucleosome binding, our research opens new avenues for investigating gene-expression regulation and perhaps developing therapeutic strategies to module gene expression in diseases associated with p53-family dysfunction.

## MATERIAL AND METHODS

### Pioneer-seq library design

The nucleosomes in the Pioneer-seq library were based on the widely used Widom-601 nucleosome-positioning sequence (Flaus 2011; Yu and Buck 2020). To ensure the Widom-601 sequence had no preexisting transcription-factor binding sites, FIMO (Grant et al. 2011) and JASPAR (Rauluseviciute et al. 2023) were used to scan this sequence for the occurrence of transcription-factor binding sites. Matching bases detected by FIMO (*p*-value < 0.001) were modified to remove detected transcription-factor binding sites.

For each of the transcription-factor binding sites included in the Pioneer-seq library, a set of Widom-601 sequences was designed, each containing a single copy of this transcription-factor binding site positioned at one of the 146 base-pair positions of the Widom-601 nucleosome core particle or at one of several linker positions.

### Pioneer-seq-nucleosome-library assembly

All library nucleosome sequences were flanked by primer sequences to generate 217-base-pair sequences. The library of nucleosome sequences, totaling 7,500 unique sequences, was acquired from Agilent as a custom oligonucleotide library, which was amplified in 15 PCR cycles using Herculase II Fusion DNA polymerase in a 100-μl reaction mixtures (1x Herculase II reaction buffer, 1 mM dNTPs, 200 pM Agilent library, 250 nM forward and reverse primers). In a typical Pioneer-seq experiment, the DNA obtained from 11 reactions was purified with a QIAquick PCR purification kit (cat. no. 28104; Qiagen), quantified with a NanoDrop spectrometer, and then visualized with a 2%-agarose ethidium-bromide gel.

Nucleosomes were generated by incubating H2A/H2B dimers and H3.1/H4 tetramers with the library DNA at a histone/DNA ratio of 1/1.5 (in a solution containing 10 mM dithiothreitol (DTT) and 1.8 M NaCl) for 30 min at room temperature. Nucleosomes were then transferred to a Slide-A-Lyzer MINI dialysis unit (10,00 MWCO; cat. no. 69750; Thermo Scientific) for dialysis performed with 1.2 ml dialysis buffers at 4°C in 1.0 M NaCl for 2 h, 0.8 M NaCl for 2 h, 0.6 NaCl for 2 h, and TE buffer (pH 8.0) overnight. After dialysis, nucleosomes were transferred to a clean 1.5-ml tube pretreated with 0.3 mg/ml bovine serum albumin (BSA), and nucleosome formation was confirmed via 4%-native-polyacrylamide-gel electrophoresis. Free DNA was then removed from nucleosomes via a 7–20%-sucrose gradient. Purified nucleosomes were then quantified via qPCR and stored at 4 °C for up to 1 month.

### Nucleosome-binding assay followed by gel-shift assay

Protein-nucleosome binding assays were performed by incubating the abovementioned purified Pioneer-seq nucleosome library with human TP53α (Abcam ab84768), TAp63α (Origene TP710041), or TAp73α (Origene TP320864) in 7 μl DNA-binding buffer (10 mM Tris-Cl (pH 7.5), 50 nM NaCl, 1 mM DTT, 0.25 mg/ml BSA, 2 mM MgCl_2_, 0.025% Nonidet P-40, and 5% glycerol) for 10 minutes on ice and then 30 minutes at room temperature. Increasing concentrations of transcription factor (0, 15, 30, 60, 120, and 240 nM) were added to 30 nM purified nucleosome library. p53-family-bound and -unbound nucleosomes were separated via gel-shift assays using 4%-native-polyacrylamide gels (acrylamide/bisacrylamide, 29:1 (w/w), 7 × 10 cm) in 0.5× Tris-borate-EDTA buffer at 100 V at 4 °C. Gel-shift assays were initially done using a wide range of p53-family-protein concentrations to determine the optimal protein amount and to ensure a shifted band is observed on the gel (**Supplemental Fig. 1**).

### DNA isolation and purification

After staining the abovementioned gel-shift assays with SYBR green (Lonza), all visible gel bands, as well as the corresponding invisible gel bands in the other lanes, were excised from the gel. Excised gel bands were then immersed in diffusion buffer (0.5 M ammonium acetate, 10 mM magnesium acetate, 1 mM EDTA (pH 8.0), 0.1% SDS) at 50 °C overnight. The gel-band-diffusion-buffer immersion was then filtered through glass wool to remove polyacrylamide, and DNA from the resultant supernatant was then purified with a QIAquick gel extraction kit (cat. no. 28704; Qiagen).

### Library construction and sequencing

Illumina sequencing libraries were prepared using a two-step PCR method. In the first step, DNA was amplified using four sets of primers (**Supplemental Table 2**) designed to offset reads and dephase the libraries during sequencing (Handelmann et al. 2023). The number of PCR cycles used in this step was determined by the DNA concentration of each sample, as measured by qPCR. In the second step, each sample was indexed using Nextera dual indices (Nextera XT index primer 1 (N7xx) and Nextera XT index primer 2 (S5xx)). After each PCR step, the reaction mixtures were cleaned up with AMPure XP beads. The concentration of each sample was then determined using the Invitrogen Quant-iT dsDNA assay kit, and equal amounts of DNA from each sample were pooled for paired-end sequencing on an Illumina NextSeq 2x150. Sequencing and quality control were performed at the University at Buffalo Genomics and Bioinformatics Core.

### Pioneer-seq analysis

FASTQ files of NextSeq reads were processed with an automated Snakemake (Koster and Rahmann 2018) pipeline of applications to refine and identify the sequences present in the sample pool. The 3’ ends of FASTQ reads with low quality scores were removed using Cutadapt (Martin 2011) with a quality cutoff of 30 (-q 30). Using Vsearch (Rognes et al. 2016), forward and reverse FASTQ reads were then merged (--fastq_mergepairs) if they shared at least 20 overlapping nucleotides (--fastq_minovlen 20) and had no more than two mismatched nucleotides between them (--fastq_maxdiffs 2). Primer sequences present at the ends of FASTQ reads were then removed using Cutadapt. FASTQ reads over 220 or under 174 nucleotides in length (--maximum-length 220 --minimum-length 174) were then filtered out using Cutadapt. Using FASTX-Toolkit (Gordon A. 2010), FASTQ reads were then converted to FASTA reads (FASTQ-to-FASTA). Then, using Vsearch (--dbmatched), each FASTQ read was mapped to a sequence in a database of the 7,500 nucleosome sequences in the Pioneer-seq library if the FASTQ read and library sequence had alignment lengths of at least 150 nucleotides (--mincols 150), had at least 98.5% similarity (- -id 0.985), and were the query and database sequence pairing with the highest percentage of identity (--top_hits_only).

The relative binding affinity of TP53, TP63, and TP73 to each of the 7,500 Pioneer-seq-library nucleosomes was calculated relative to control Pioneer-seq-library nucleosomes (i.e., nucleosomes with p53-family-nonspecific binding sites—e.g., binding sites of FoxA1, Klf4, Stat3, etc.):

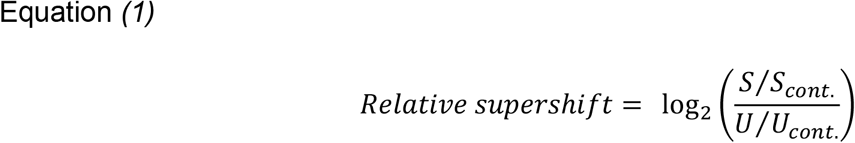

where, for a given nucleosome in the library, *S* is the number of reads of this given nucleosome in a shifted gel band, *S*_*cont*._ is the average number of reads of library nucleosomes with p53-family-nonspecific binding sites in the same shifted gel band, *U* is the number of reads of this given nucleosome in the unshifted gel band from the lane of the gel to which no transcription factor was added, and *U*_*cont*._ be the average number of reads of library nucleosomes with p53-family-nonspecific binding sites in the same unshifted gel band.

Pioneer-seq experiments were initially performed using multiple concentrations of p53-family protein in case of differences in inherent binding affinities and protein purity. The p53-family concentrations that enabled the highest p53-family-binding-site-specific binding—the concentrations used throughout this paper—were as follows: 15 nM TP53, 60 nM TP63, and 60 nM TP73.

### Nucleosome-binding assay using Cy5-labelled nucleosomes

Nucleosomal DNA labeled with the fluorescent cyanine dye Cy5 on its 5’ and 3’ ends was formed into purified nucleosomes as described above. Nucleosomes (30 nM) were incubated with increasing amounts of p53-family protein (0, 15, 30, 60, 120, 240 nM) in 7 μl DNA-binding buffer on ice for 10 min and then at room temperature for 30 min. p53-family-bound and -unbound nucleosomes were then separated via gel-shift assays using 4%-native-polyacrylamide gels in 0.5× Tris-borate-EDTA buffer at 100 V at 4 °C. After the gel shift assays, the nucleosomes were visualized and quantified via their Cy5 labels using a ChemiDoc MP imaging system. The intensity of the Cy5 fluorescence was directly proportional to the amount of nucleosomes present, enabling the quantification of the percentage of nucleosome bound.

### MNase-digestion of Pioneer-seq nucleosome library

MNase was used to measure the accessibility of nucleosomes as previously described (Handelmann et al. 2023). The Pioneer-seq nucleosome library (30 nM) was digested by MNase (0.05 U/μl) in nuclease digestion buffer (10 mM Tris-HCL (pH 8.0), 2 mM CaCl_2_) at 37 °C for 0, 5, or 10 minutes; digestion was stopped with 2% SDS and 40 mM EDTA. Each sample was then incubated with proteinase K (16 μg) for 1 h at 55 °C. DNA from each sample was then purified and concentrated with the QIAquick PCR purification kit. Concentrations of DNA in each sample were then determined with the Invitrogen Quant-iT dsDNA assay kit and equalized. Illumina sequencing libraries were generated using an NEBNext Ultra II DNA library prep kit. Individual samples were multiplexed for paired-end sequencing on an Illumina MiSeq 2x150. Sequencing and quality control were performed at the University at Buffalo Genomics and Bioinformatics Core.

FASTQ files of MiSeq reads were quality-filtered (*q* > 30) and adapter-trimmed using Cutadapt. The filtered and trimmed reads were then merged and mapped to a database of the 7,500 nucleosome sequences in the Pioneer-seq library using Vsearch. Based on the number of reads and the end positions of these reads, MNase-digestion scores were calculated for each base pair of library nucleosomes as the ratio of the number of reads covering that base pair to the total number of reads mapped to the respective library nucleosome.

### Modeling Pioneer-seq-library nucleosomes and measuring solvent-accessible surface area

Structural models for nucleosomes in the Pioneer-seq library were generated using version 1.7 of ChimeraX (Goddard et al. 2018) and based on the crystal structure of a Widom-601 nucleosome (PDB accession no.: 4QLC). The linker DNA on one side of the Widom-601 crystal structure was extended to match that of the nucleosomes in the Pioneer-seq library.

The solvent-accessible surface area (SASA) of the two CATGs within the high-affinity p53-family binding site was calculated using the ‘measure sasa’ ChimeraX command with a ‘probeRadius’ of 0.9 Å.

## DATA AVAILABILITY

All Pioneer-seq results from this study are available in the Sequence Read Archive under the accession number PRJNA1048449.

## ACKNOWLEDGEMENTS

This study was supported by the National Institute of General Medical Sciences [R01GM132199] to MJB. We thank the UB Genomics and Bioinformatics Core for next-generation sequencing services.

